# IIHP: Intelligent Incident Hypertension Prediction in Obstructive Sleep Apnea

**DOI:** 10.1101/2023.12.13.571552

**Authors:** Omid Halimi Milani, Ahmet Enis Cetin, Bharati Prasad

## Abstract

The Obstructive sleep apnea (OSA) increases the risk of hypertension, mainly attributed to intermittent hypoxia and sleep fragmentation. Given the multifaceted pathogenesis of hypertension, accurately predicting incident hypertension in individuals with OSA has posed a considerable challenge. In this study, we leveraged Machine Learning (ML) techniques to develop a predictive model for incident hypertension up to five years after OSA diagnosis by polysomnography. We used data from the Sleep Heart Health Study (SHHS), which included 4,797 participants diagnosed with OSA. After excluding those with pre-existing hypertension and Apnea Hypopnea Index (AHI) values below 21 per hour, we had 671 participants with five-year follow-up data. We adopted two distinct methodologies. We first implemented adaptive convolution layers to extract features from the signals and combined them into a 2D array. The 2D array was further processed by a 2D pre-trained neural network to take advantage of transfer learning. Subsequently, we delved into feature extraction from full-length signals across various temporal frames, resulting in a 2D feature array. We studied the use of various 2D networks such as MobileNet, EfficientNet, and a family of RESNETs. The best algorithm achieved an average area under the curve of 72%. These results suggest a promising approach for predicting the risk of incident hypertension in individuals with OSA, providing tools for practice and public health initiatives.

## 1 Introduction

Obstructive sleep apnea (OSA) is a condition characterized by intermittent hypoxia and sleep fragmentation, which propagates hypertension via mechanisms such as sympathetic activation and inflammation. OSA also raises the likelihood of developing hypertension during nighttime. [1-4]. Notably, specific patterns of apnea, such as Rapid Eye Movement (REM) sleep-related OSA, could contribute to OSA-associated hypertension [5]. Additional sleep-related factors, like reduced slow-wave sleep (SWS) and short sleep duration, are associated with hypertension, independent of OSA [6, 7]. Ren and colleagues conducted a study examining the association, between sleep duration, obstructive sleep apnea (OSA) and hypertension in a group of 7,107 OSA patients and 1,118 primary snorers. The findings from polysomnography indicated that individuals who slept for 5 to 6 hours had a 45% risk of developing hypertension while those who slept, than 5 hours had an 80% increased risk. These results were independent of any factors that could potentially influence the outcome. [8].

Around 30% of people, in the United States are affected by hypertension, which can lead to dysfunction in organs. Treating obstructive sleep apnea (OSA) has been shown to lower the risk of developing hypertension. [9]. However, accurately predicting the onset of hypertension in individuals with OSA remained a challenge due to the complex pathogenesis of hypertension.

In recent years, there has been a growing recognition of the limitations associated with conventional manual sleep stage scoring, which simplifies the analysis of electroencephalogram (EEG) temporospectral and frequency domains. This scoring method is inherently subjective and can lead to variations between different scorers due to the application of visual-based rules [10]. To reduce this variability, researchers used power spectral density (PSD) analysis of EEG signals, which objectively examines sleep EEG microarchitecture. This approach enables the decomposition of EEG brain waves across various power frequency bands, ranging from slow wave activity (delta EEG power, 1–4 Hz) to fast-frequency activity (beta EEG power, 18–30 Hz), achieved through implementing fast Fourier transform algorithms. At a microarchitecture level, slow wave sleep (SWS) is characterized by high delta power, indicative of deep sleep. Numerous studies have suggested that quantitative EEG analysis may yield more sensitive biomarkers for adverse health outcomes in OSA compared to traditional sleep scoring methods [11-14]. Other features that also can be useful to study when working with EEG signals [15]. However, there is a paucity of research investigating the relationship between sleep microarchitecture and the future development of hypertension in OSA. Berger et al. recently reported that low delta power in non-REM sleep is independently associated with the risk of developing hypertension, confirming the previously noted relationship of SWS with incident hypertension [16]. Another study relied on complex feature engineering to examine 27 unselected clinical predictors, encompassing demographic characteristics, lifestyle behaviors, and OSA severity (apnea-hypopnea index; AHI, and oximetry indices) [17].

Ruitong et al. adopted a pulmonary physiology-based approach to predicting the onset of hypertension by including pulmonary function measurements and polysomnography-derived indices using a penalized regression and Elastic Net model [18]. A recent study introduced cSP (sleep and pulmonary) phenotypes, which combines spirometry and overnight polysomnography measures to predict hypertension occurrence in the Sleep Heart Health Study (SHHS). The researchers employed penalized regression to select features and estimate effect size simultaneously[11]. The SHHS dataset encompasses a variety of physiological signals, including sleep EEG and electrooculogram (EOG), electrocardiogram (ECG), electromyogram (EMG), ventilatory effort, nasal airflow, photoplethysmography-derived oximetry, snoring, and body position. Employing rigorous signal processing techniques, such as filtering, segmentation, and feature extraction, our analysis aimed to unravel the intricate patterns embedded within these signals [19, 20].

This study leveraged deep Machine Learning (ML) techniques to develop a predictive model for incident hypertension up to five years after OSA diagnosis by polysomnography. The novel ML-based method utilized polysomnography signal conversion and a composite deep CNN model to transform the polysomnography signals into image datasets to predict incident hypertension. During the training process, additional static information for each patient was incorporated, including age, sex, race, body mass index, baseline systolic, and diastolic blood pressure. Our study aimed to accelerate precision medicine in managing hypertension and preventive healthcare by integrating polysomnography and clinical data analytics. This research could equip healthcare professionals with tools to address the significant health burden posed by OSA and hypertension [21].

We hypothesized that a comprehensive approach involving the simultaneous input of time-series physiological signals measuring sleep (EEG), ventilatory impairment and hypoxia, and cardiac autonomic dysregulation (electrocardiogram and photoplethysmography-derived heart rate variability and pulse transit time) preserved the temporal correlations between multiple physiological perturbations in OSA and could provide a robust prediction of incident hypertension in OSA. These signals were included in polysomnography, a widely available diagnostic test for OSA. Thus, we extracted multiple features from the polysomnography signals in the SHHS participants with moderate to severe OSA. [19, 20, 22, 23]. Before feature extraction, rigorous preprocessing steps were employed, involving removing artifacts and applying a bandpass filter to eliminate any remaining noise. Next, we explored whether the addition of clinical factors that independently affected the risk of hypertension (age, sex, race, body mass index, and baseline systolic and diastolic blood pressure) improved the prediction performance of the ML model.

The paper advocates using timing interval windowing as an effective approach to feature extraction. This technique enabled the creation of a structured 2D array, facilitating the application of transfer learning. Additionally, a model fine tuning process was applied to enhance its adaptability to the specific characteristics of the sleep data. The paper introduces a novel adaptive feature extractor, designed from the ground up. This extractor was meticulously crafted to adapt to the intricacies of the sleep data, ensuring a comprehensive and accurate representation of the underlying patterns. We analyzed the signal using networks. Initially, we abstracted details from the study, constructed an image based on these details, and customized the initial EfficientNet model, which was pretrained, for the task by adjusting its layers. To overcome limitations posed by the number of features and limited memory capacity, a 32-bit model was utilized. Our second approach focused on using networks (CNNs) to extract features from the signals. We created a 2D array that represented segments of the signals and their associated features. These features included counts of arousal and respiratory events, as well as statistical measures like mean, standard deviation, skewness, and kurtosis for heart rate variability analysis. After exploring timing intervals, we determined that a window length of 10 minutes showed promising results.

The proposed methodology underwent a rigorous evaluation through a 10-fold cross-validation approach to examine the model’s generalizability. The results were subsequently summarized, comparing models and methods to cutting-edge approaches.

In the following sections, we explain our methodology, present our findings, and discuss the implications of our research within the context of practice and public health initiatives. Section 2 detailed our methodology, covering feature extraction, machine learning models, and innovative techniques. Section 3 briefly summarizes results, comparing methods with references. Section 4 and 5 are discussion and conclusion respectively. The conclusion section discusses the implications of this study and future research.

## 2 Methods

In this study, we explored two methodologies for predicting incident hypertension in individuals with moderate to severe OSA. One notable enhancement to our approach involved the integration of a convolution layer designed to automatically extract features from each signal. This layer dynamically adjusted itself during training, facilitating the automatic extraction of features and identification of crucial patterns within the data. However, it came with a computational expense, leading us to modify the computation to 32-bit, albeit extending the training time to 10 days. In an alternative approach, we devised a system to extract features directly from the signals. The extracted features from the entire signals were then converted into a 2D array. Each row in the array represented the feature representation of a signal, enabling the capture of temporal information from diverse signals through the application of convolution kernels. This method was not only easy to train but also faster, allowing our server to complete the training process within 4 hours. Overall, our findings indicated that the second method, with its efficiency and superior model performance, stood out among the experiments. Nevertheless, with the utilization of faster and more capable servers, coupled with an expanded dataset, there is potential for reconsidering and refining the other two methods. The implementation details are listed in the following sections. Then, we highlighted the details of the findings and reported the performance comparison with the state-of-the-art methods.

Prior to conducting any data analysis, we implemented a preprocessing step for each signal. Specifically, a portion of the SaO2 Signal, along with OX, underwent processing in Module 1. Notably, the presence of numerous sparks due to power outages necessitated their removal using the Spark Remover filter. To address this issue, we employed Module 1 and Filter 1 as depicted in Table 1.

**Table 1.**
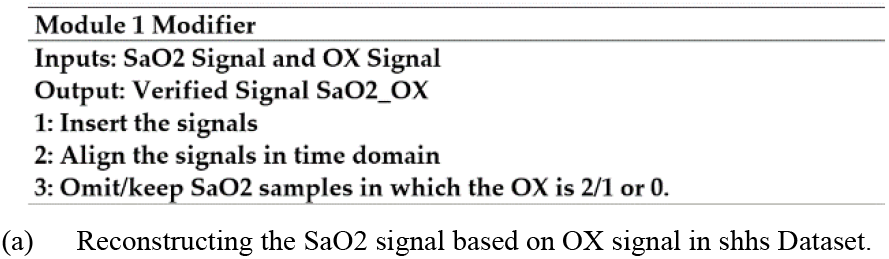
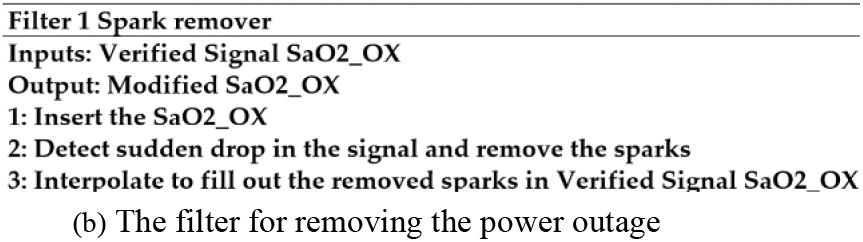
As indicated in the dataset, we can utilize OX for signal validation in each second, and it is essential to remove sparks, as described in Module 1 and Filter 1 respectively.

### 2.1 Intelligent Signal-Based Method for Predicting Incident Hypertension from Obstructive Sleep Apnea

The intelligent signal-based method involved extracting features from signals using convolutional layers to construct an image from the derived features. These features were then inputted into the pre-trained EfficientNet, with fine-tuning applied to the first convolutional layers and the final classifier layer. The overall methodology was depicted in Figure 2, where the red boxes represented the kernels, scaled according to the sampling frequencies of the signals. Following the application of convolution to each signal, the results from different signals were consolidated and passed through a converter, constituting another convolutional layer that standardized the input for the EfficientNet. Consequently, the 11 channels were condensed to 3 channels before being fed into the EfficientNet. All CNN models were trained, in addition to the EfficientNet classifier.

The signals’ names in this study were based on the SHHS study. The SHHS signal comprised vital parameters for sleep analysis. “sao2_OX” monitored blood oxygen levels, EEG tracked brain activity, and EMG recorded muscle activity. EEG (sec) provided additional brain activity details. EOG(L) and EOG(R) monitored eye movements. THOR and ABDO measured respiratory effort, POSITION recorded body posture, and NEW AIR tracked fresh air introduction. This diverse set of signals allowed for a detailed assessment of sleep patterns, respiratory functions, and positional variables, contributing to a comprehensive understanding of sleep health [19, 20].

Figure 1 depicted the structure of the methodology and how the data fed into our model after different artifacts from the signals were removed. This approach faced computational challenges when applied to a single patient’s data, given the extended signal duration (around 9 hours) with approximately 4 million data points in the ECG signal alone. The combination of signals resulted in a high feature count, exacerbated by a limited dataset unsuitable for full model training. To overcome these constraints, a proposed solution involved a novel approach to feature extraction through windowing.

**Fig. 1.**
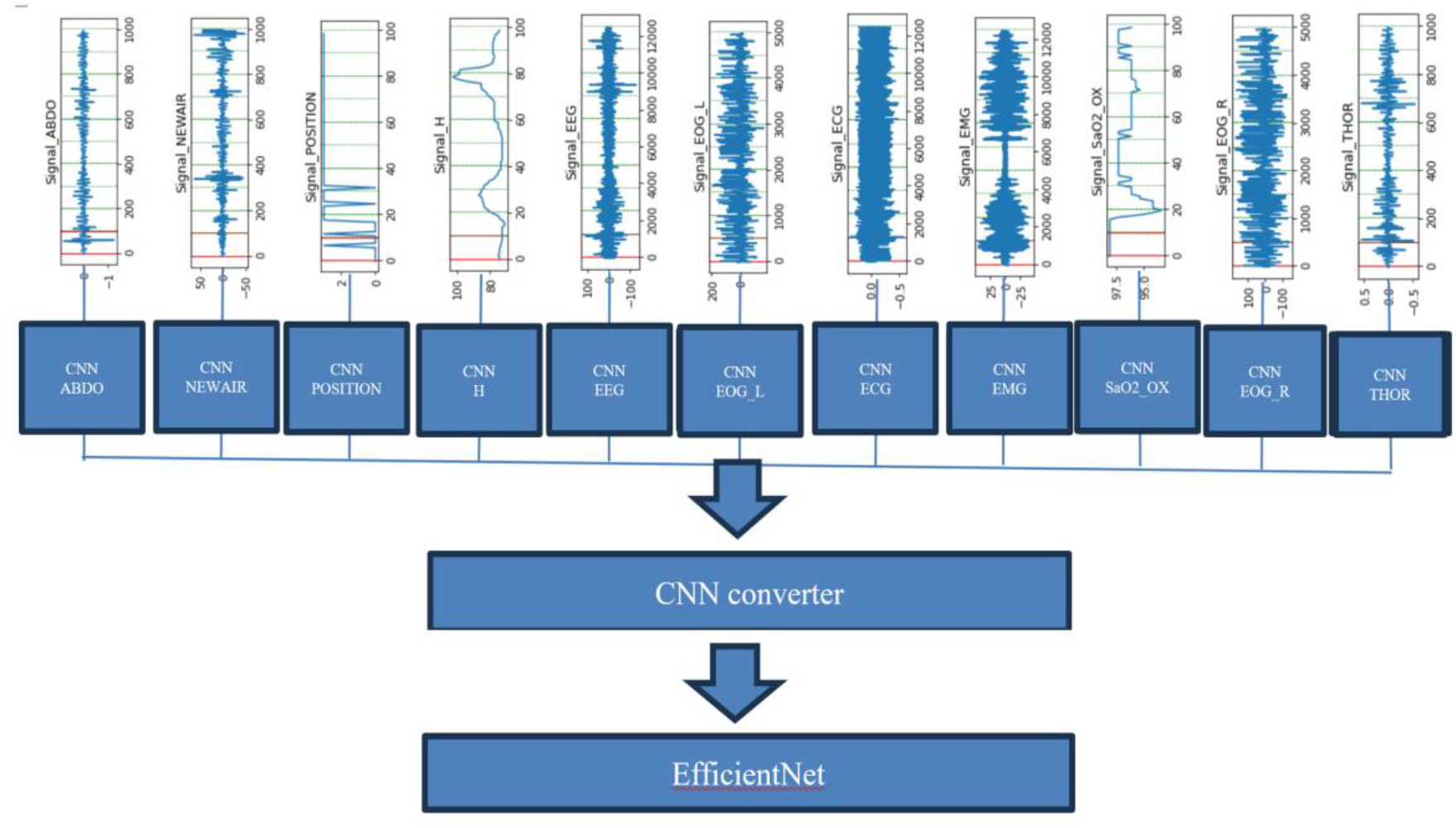
Feature extraction from signal windows using Convolutional Neural Networks (CNN). The CNN architecture enhances pattern recognition and understanding.

### 2.2 Intelligent Feature-Based Method for Predicting Incident Hypertension from Obstructive Sleep Apnea

The occurrence of hypertension appears to be non-trivial to pinpoint within the signals. Therefore, we adopted a holistic approach, considering the entire signal rather than an event-based strategies. In this method, we applied various windowing sizes to the signals and extracted the relevant features from each segment using different CNNs processing individual signals. These features were then organized into a 2D array, where each column represented a segment of the signal along with its corresponding features. The features encompassed the frequency of arousal events, respiratory events, heart rate variability, as well as statistical measures like minimum, standard deviation, mean, skewness, kurtosis, maximum, and median derived from 11 distinct signals.

Several statistical features extracted from the Signals. The mean of a dataset in equation 1, denoting the average value, was calculated using the following formula where 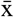 was the mean, n was the number of data points, and x_i_ are the individual data points.

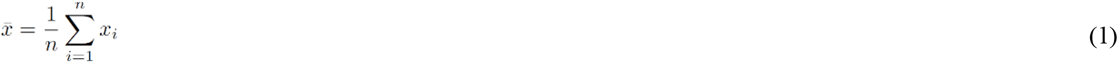

The standard deviation, as shown in Equation 2, measured the spread or dispersion of the data points around the mean. [24]. It was calculated as:

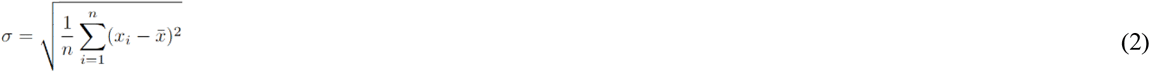

The symbol σ represents the deviation n stands for the number of data points 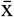 represents the mean. Xi refers to each individual data point. Skewness is a measure of how asymmetrical the probability distribution of a valued variable is. A positive skewness, calculated using equation 3 as mentioned in [25] suggests a tail, on the side whereas a negative skewness indicates a longer tail, on the left side.

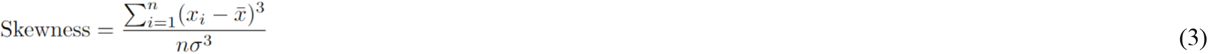

Kurtosis, as calculated in the equation, measures the ‘tailedness’ of a probability distribution. It describes the shape of the distribution’s tails relative to the normal distribution. [25].

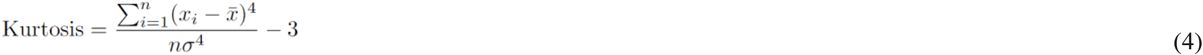

Another crucial feature involved heart rate variability (HRV), a parameter integral to the assessment of cardiac health and autonomic nervous system function. HRV was determined by detecting peaks and variations in the time intervals between successive heartbeats and corresponding HRV features calculated as showed in Table 2. This metric provided valuable insights into the adaptability and resilience of the cardiovascular system, reflecting the dynamic interplay between the sympathetic and parasympathetic branches of the autonomic nervous system [26, 27].

**Table 2.**
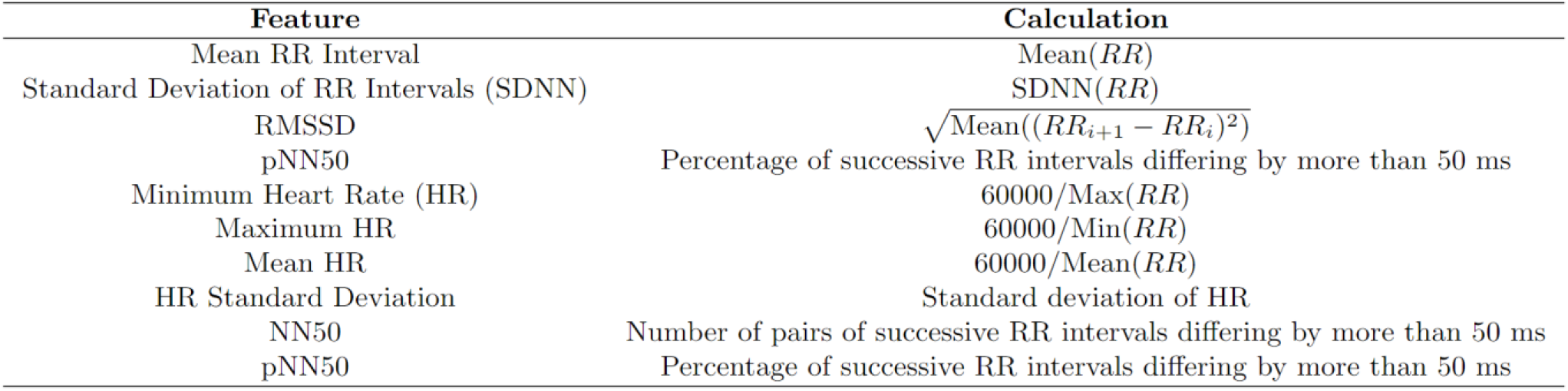
The Heart Rate Variability (HRV) Calculation Algorithm is a systematic process designed to extract essential features from raw Electrocardiogram (ECG) signals.

| Feature | Calculation |
| --- | --- |
| Mean RR Interval | Mean(RR) |
| Standard Deviation of RR Intervals (SDNN) | SDNN(RR) |
| RMSSD | $\sqrt{\text{Mean}((RR_{i+1} - RR_i)^2)}$ |
| pNN50 | Percentage of successive RR intervals differing by more than 50 ms |
| Minimum Heart Rate (HR) | 60000/Max(RR) |
| Maximum HR | 60000/Min(RR) |
| Mean HR | 60000/Mean(RR) |
| HR Standard Deviation | Standard deviation of HR |
| NN50 | Number of pairs of successive RR intervals differing by more than 50 ms |
| pNN50 | Percentage of successive RR intervals differing by more than 50 ms |

**Table 3.**
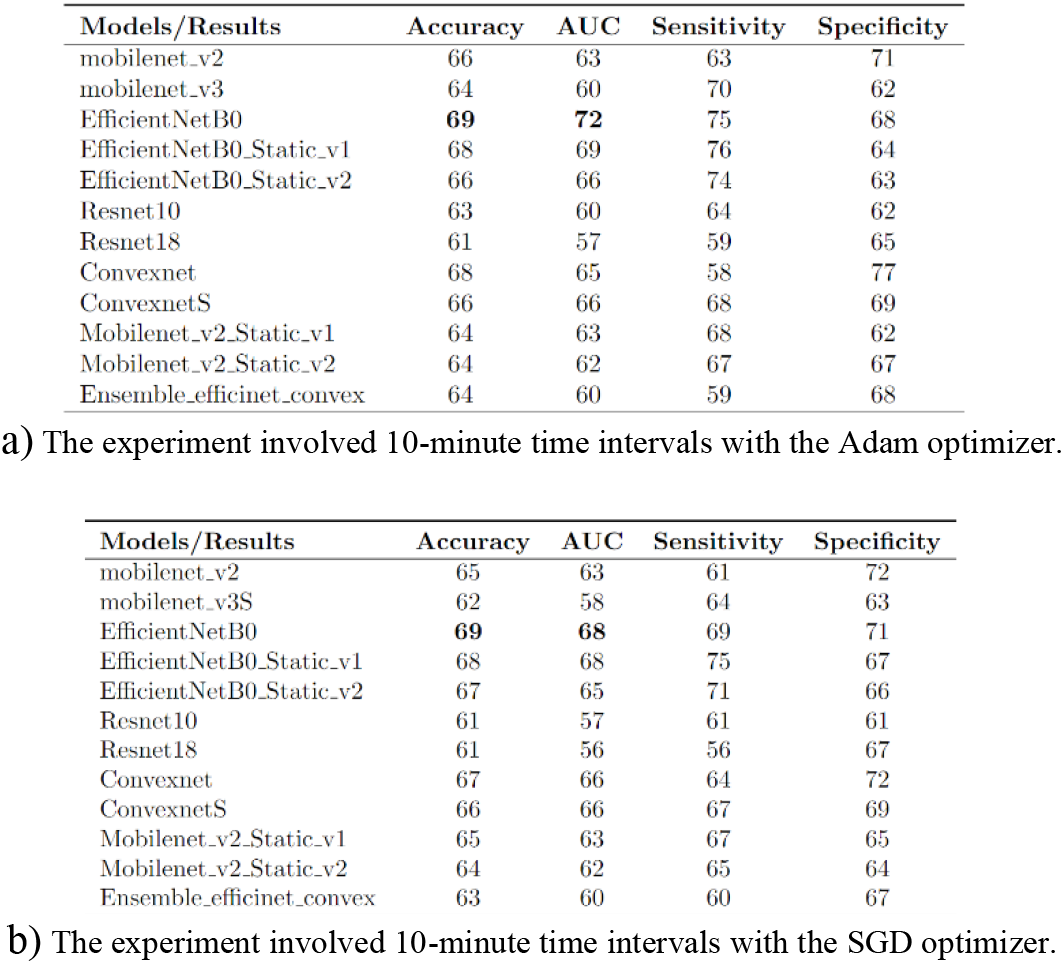
The structure of different transfer learning models were examined through various experiments. Experiments were conducted on the images (2D arrays) generated from the extracted features: a) 2D arrays derived from the extracted features with 10-minute intervals and the Adam optimizer. b) 2D arrays derived from the extracted features with 10-minute intervals and the SGD optimizer.

**Table 4.**
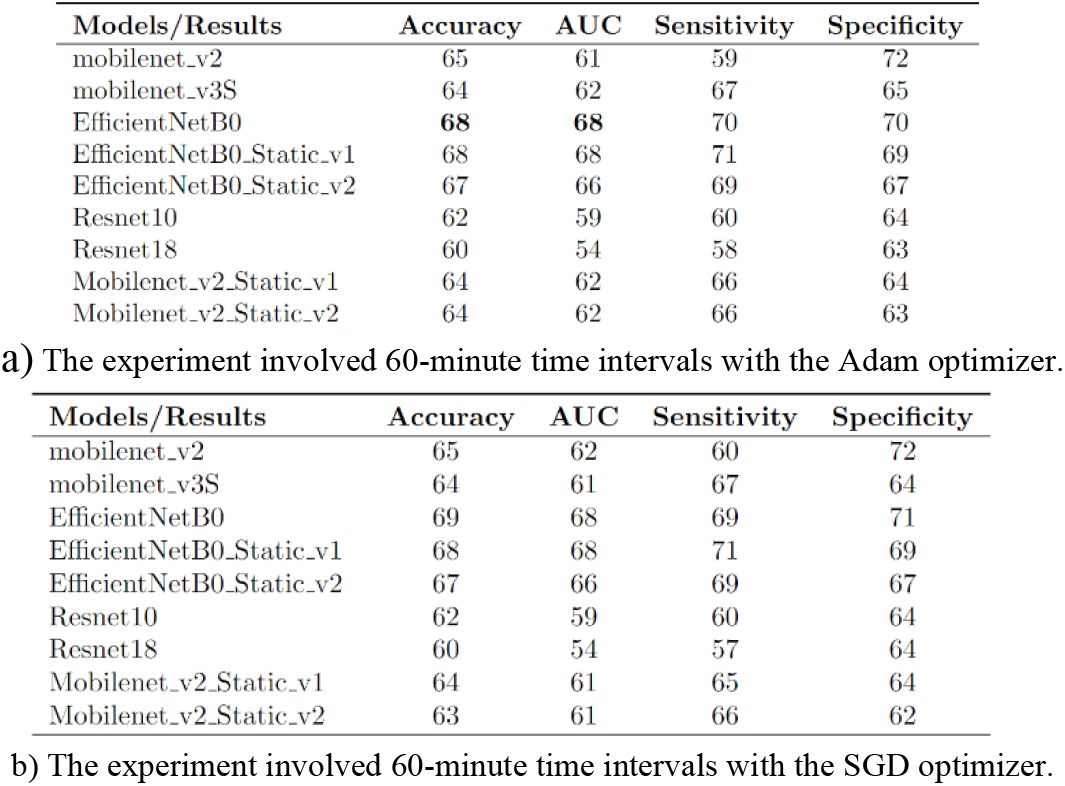
a) 2D array derived from the extracted features with 60-minute intervals and Adam optimizer. b) 2D array derived from the extracted features with 60-minute intervals and SGD optimizer.

Figure 2 outlined the steps involved in constructing the given dataset. Depending on the window timing, different features were extracted from the signals. These included minimum, standard deviation, average, skewness, kurtosis, maximum, and median within the T-minute interval of the signal. Additionally, the count of respiratory and arousal events within that T-minute interval was recorded. Furthermore, heart rate variability was derived from the ECG signals. Together, these components constituted the first column in the 2D array. Subsequent columns followed the same pattern, with the window shifting to the right with a stride of T minutes.

**Fig. 2.**
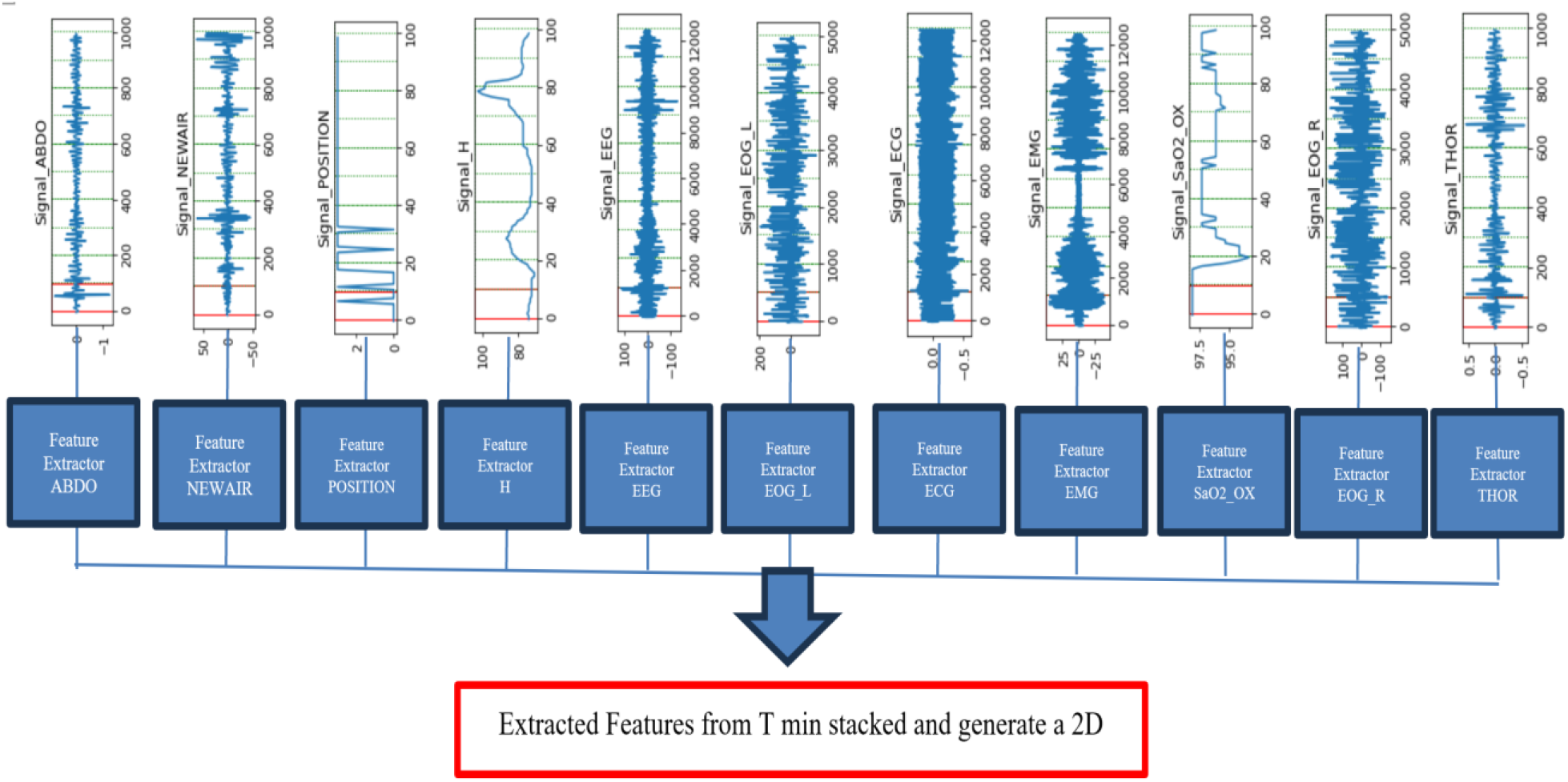
Feature extraction from signal windows using various statistical metrics involved computing features such as maximum, minimum, average, standard deviation, kurtosis, skewness, heart rate variability, number of arousal events, and number of respiratory events. These diverse features contributed to comprehensive pattern recognition and understanding within the signal processing framework.

The potential of employing Finite Impulse Response (FIR) and Infinite Impulse Response (IIR) filters for data augmentation was also explored. These filters were applied to the signal prior to extracting features from them. Different orders of 4, 6, and 8 were applied to the signals for FIR filters, and the impulse response of an FIR filter was a sequence of finite duration. Different orders of 18, 20, and 22 were applied to the signals for IIR filters [28].

The Convolutional Neural Network (CNN) was a widely adopted and highly efficient trainable model for tasks related to image classification. Its effectiveness in categorizing images into specific classes had made it an indispensable tool in various computer vision applications. By utilizing 2-D convolution operations, CNNs were able to extract meaningful information from images. These convolution layers acted as feature extractors, capturing crucial patterns, textures, and structures present in the input images.

In recent years, significant progress has been made in the field of computer vision, leading to the development of numerous CNN architectures, each aiming to achieve higher accuracy on large benchmark datasets. Among these architectures, ResNet stands out for effectively addressing the vanishing gradient problem in deep CNNs. Through the introduction of residual connections, information flows directly through the network, allowing ResNet to achieve greater depth, expressiveness, and improved image classification performance.

Another notable advancement in CNN architectures was MobileNet. These models were characterized by their lightweight and resource-efficient nature, making them well-suited for deployment on less powerful devices such as mobile phones and embedded systems. They utilized depthwise separable convolutions, which significantly reduced parameters and computations compared to traditional convolutions. This resulted in faster inference times and optimal performance for real-time applications on resource-constrained devices. [29]. Like MobileNet, ShuffleNet is a special kind of computer program for mobile phones. It helps them do things quickly without using too much power. ShuffleNet has two clever parts that work together to save power and still do things accurately [30].

EfficientNet was a revolutionary CNN architecture that merged the advantages of ResNet and MobileNet, addressing their limitations. It employed compound scaling to optimize network depth, width, and resolution simultaneously with a single scaling coefficient. This enabled a balanced trade-off between model size and accuracy, enhancing versatility and efficiency for diverse image classification tasks. The architecture integrated depthwise separable convolutions from MobileNet and residual connections from ResNet, achieving state-of-the-art performance on benchmark datasets with improved computational efficiency [31].

In this study, we primarily employed EfficientNet-B0 as our base model, comparing its results with various other CNN architectures. Our proposed algorithm built upon EfficientNet as the baseline, incorporating comparisons with other models. The input image shape consisted of a 2D array where rows represented features, and columns denoted the specified window time, set at 10 minutes. This 2D array was replicated and input as different channels to EfficientNet, with all CNN model classifiers being fine-tuned.

The static data was also experimented with in different models. The way the static data was included in the decision-making process varied, as described in Figure 3. One approach involved adding the static data to the final layer of the pretrained models and fine-tuning all the weights together to satisfy the desired output. Another approach involved adding another fully connected layer and concatenating the static data in the second layer, followed by fine-tuning all the weights together. In an effort to improve our abilities, we decided to combine two models: one based on EfficientNetB0 and the other using the ConvexNet architecture. However, the combination of these models did not noticeably enhance our performance. This suggests that both models may have generated predictions resulting in synergy [32, 33]

**Fig. 3.**
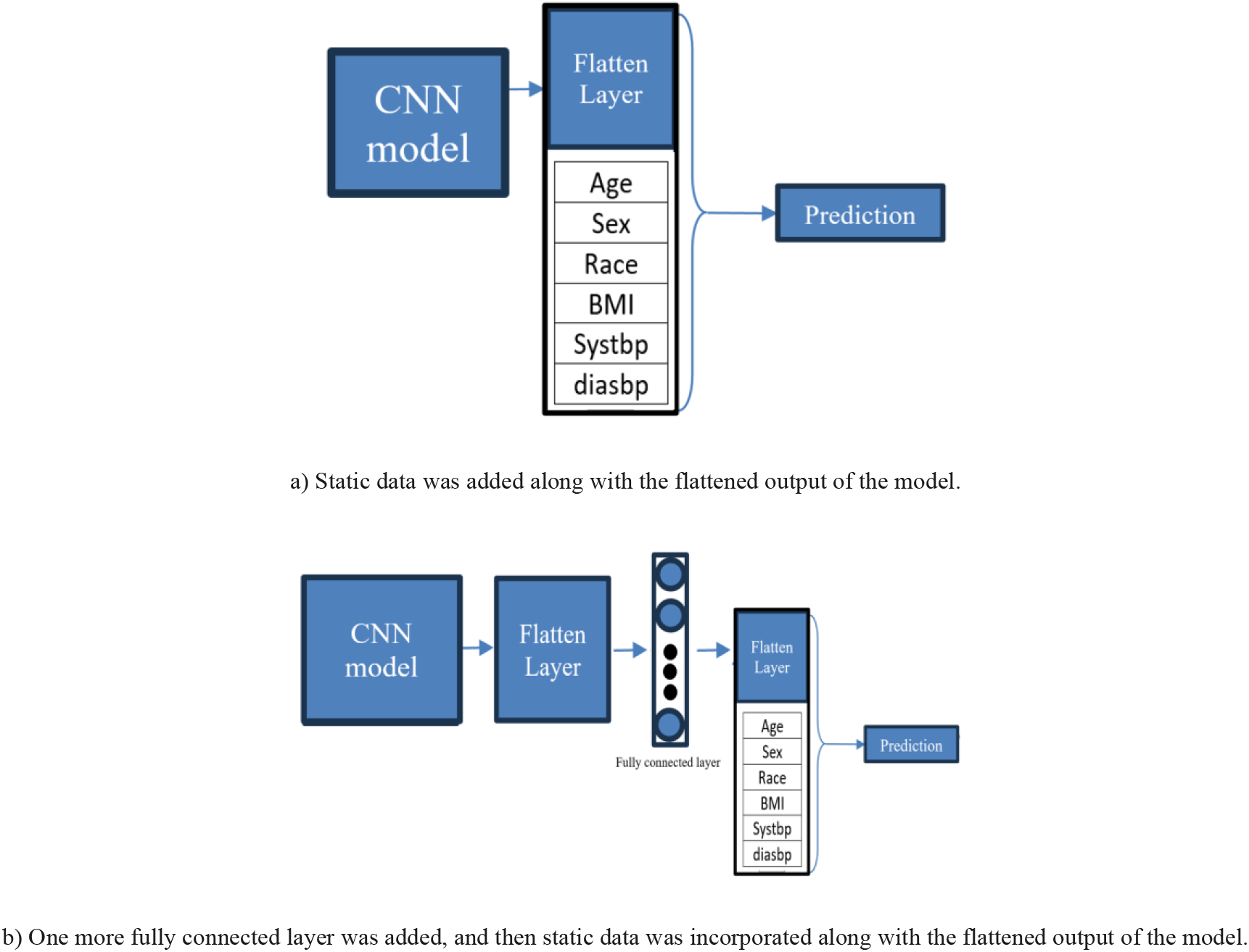
The static data, including BMI (body mass index), Sysbp (systolic blood pressure), and Diasbp (diastolic blood pressure), was added to the model. a) static_v1, b) static__v2

## 3 Results

We employed various metrics to evaluate the robustness of our approaches. To assess the model’s performance, we utilized 10-Fold cross-validation to mitigate potential biases. The process for implementing 10-fold cross-validation was outlined in Figure 4. This involved partitioning the training data into training and validation sets, with each round validating the data against the trained model. Ultimately, the results were averaged across the 10 different folds.

**Fig. 4.**
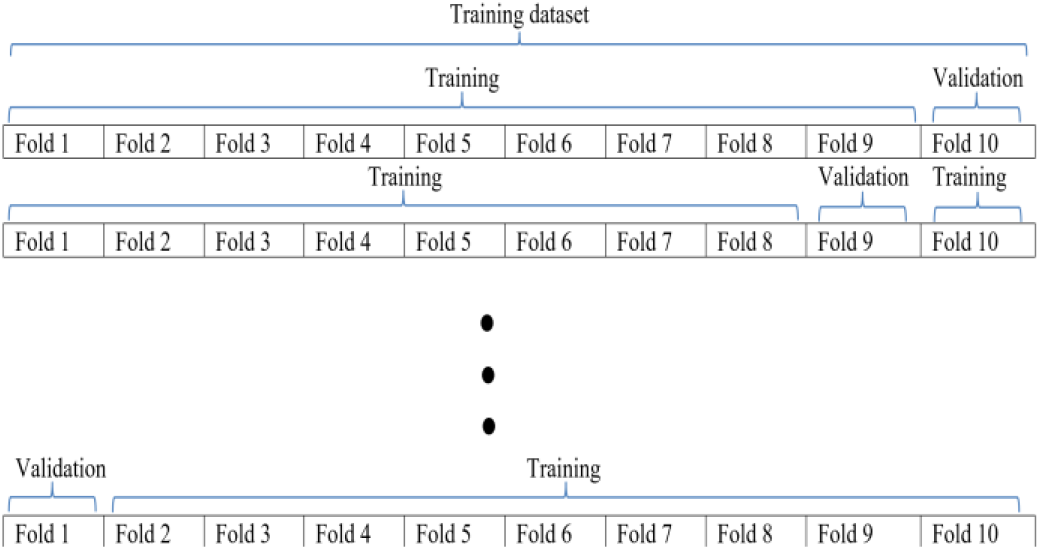
10-fold cross validation method

We used equations 7, 8 and 9 to calculate the accuracy, area, under the curve (AUC) using rate (TPR) and false positive rate (FPR) respectively. The AUC was determined by plotting TPR, against FPR at classification thresholds. It helped us measure how well the model can distinguish between classes with values indicating performance.

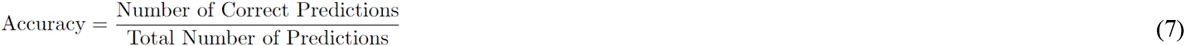

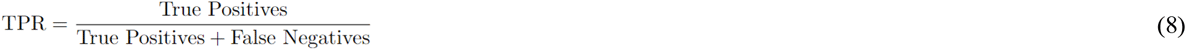

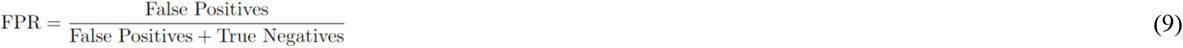

Choosing the optimal range for sensitivity and specificity, or their equivalents TPR and FPR, was crucial for our study. Sensitivity (True Positive Rate) took precedence when detecting positive cases was critical, while specificity (1 - False Positive Rate) was vital for minimizing false alarms.

In Approach 1 and 2, we utilized Convolutional Neural Networks (CNNs). We explored state-of-the-art pretrained models for transfer learning after preprocessing the data points. The input data fed into the model consisted of 2D arrays, obtained by applying varying windowing techniques on the signal proportional to the sampling frequencies. The difference between approach 1 and 2 was about their feature extraction methods. We conducted experiments with different window lengths and assessed the model’s performance under various intervals to identify the near-optimal windowing strategy. Our findings indicated that a 10-minute window length yielded the most favorable results compared to other timing intervals with the second approach. Additionally, we investigated both Adam and SGD as optimizers, coupled with schedular techniques, to explore the solution space more effectively.

In addition to the 10-minute and 60-minute intervals, we investigated various timing intervals. The results for the best models at different timing intervals were summarized in Table 5.

**Table 5.**
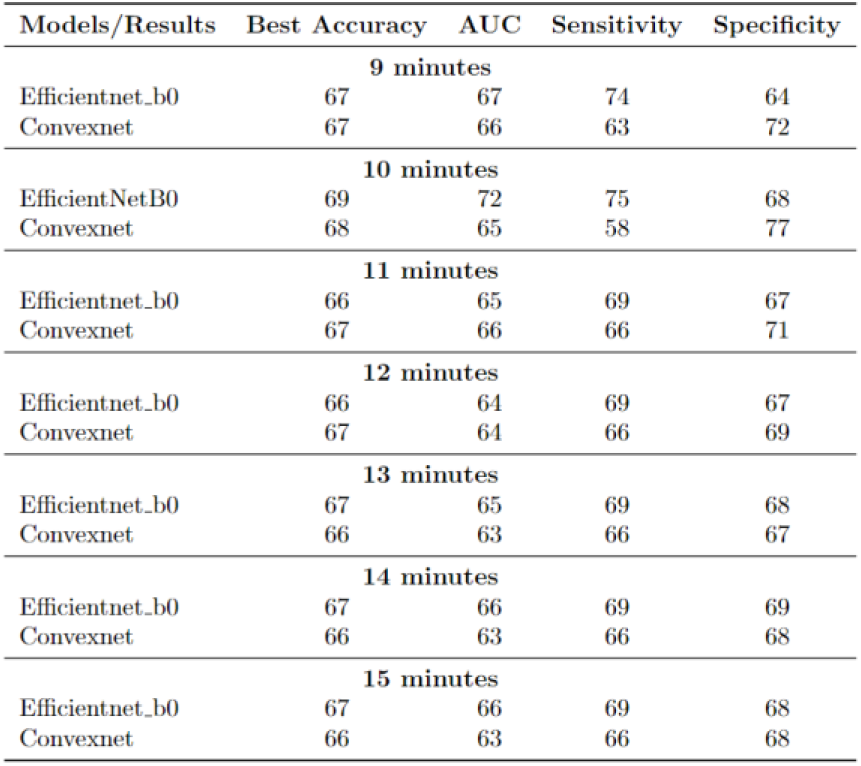
The results for different windowing on the signal for feature extraction.

| Models/Results | Best Accuracy | AUC | Sensitivity | Specificity |
| --- | --- | --- | --- | --- |
| 9 minutes |  |  |  |  |
| Efficientnet_b0 | 67 | 67 | 74 | 64 |
| Convexnet | 67 | 66 | 63 | 72 |
| 10 minutes |  |  |  |  |
| EfficientNetB0 | 69 | 72 | 75 | 68 |
| Convexnet | 68 | 65 | 58 | 77 |
| 11 minutes |  |  |  |  |
| Efficientnet_b0 | 66 | 65 | 69 | 67 |
| Convexnet | 67 | 66 | 66 | 71 |
| 12 minutes |  |  |  |  |
| Efficientnet_b0 | 66 | 64 | 69 | 67 |
| Convexnet | 67 | 64 | 66 | 69 |
| 13 minutes |  |  |  |  |
| Efficientnet_b0 | 67 | 65 | 69 | 68 |
| Convexnet | 66 | 63 | 66 | 67 |
| 14 minutes |  |  |  |  |
| Efficientnet_b0 | 67 | 66 | 69 | 69 |
| Convexnet | 66 | 63 | 66 | 68 |
| 15 minutes |  |  |  |  |
| Efficientnet_b0 | 67 | 66 | 69 | 68 |
| Convexnet | 66 | 63 | 66 | 68 |

The overall results for different models and methods, along with a comparison to the state of the art, were presented in Table 6.

**Table 6.** The results for different windowing on the signal for feature extraction were examined.

| Models/Results | Best Accuracy | AUC | Sensitivity | Specificity |
| --- | --- | --- | --- | --- |
| mobilenet_v2 | 66 | 63 | 63 | 71 |
| mobilenet_v3 | 64 | 60 | 70 | 62 |
| EfficientNetB0 | 69 | 72 | 75 | 68 |
| EfficientNetB0_Static.v1 | 68 | 68 | 71 | 69 |
| EfficientNetB0_Static.v2 | 67 | 66 | 69 | 67 |
| Resnet10 | 63 | 60 | 64 | 62 |
| Resnet18 | 61 | 57 | 59 | 65 |
| Convexnet | 68 | 65 | 58 | 77 |
| Convexnet_Static.v1 | 66 | 66 | 68 | 69 |
| Mobilenet_v2_Static.v1 | 64 | 63 | 68 | 62 |
| Mobilenet_v2_Static.v2 | 64 | 62 | 67 | 67 |
| Ensemble_efficient_convex | 64 | 60 | 59 | 68 |
| EfficientNetb0_FIR_IIR | 64 | 60 | 59 | 68 |
| cSPPSG [11] | - | 71 | - | - |
| AHI [11] | - | 67 | - | - |
| EfficientNetb0_CNN_feature | 65 | 63 | 68 | 63 |

EfficientNetB0 emerged as the top-performing model with the highest Best Accuracy (69%) and AUC (72%), showcasing its effectiveness. Other models, such as Convexnet also demonstrated competitive performance across different metrics. The table served as a comprehensive overview of the comparative performance of various models in the given study context.

## 4 Discussion

The research discussed in this paper aimed to create a model that can predict the likelihood of developing hypertension in individuals with sleep apnea (OSA) using machine learning techniques applied to data from polysomnography tests. By combining machine learning methods and comprehensive feature extraction techniques our study marks a step forward in the field of sleep medicine potentially impacting clinical practices and public health efforts.

The findings from our study show promising results in forecasting the onset of blood pressure in individuals with OSA achieving an accuracy rate of 72%. This indicates that our predictive model has the ability to effectively identify individuals who’re at risk of developing blood pressure after being diagnosed with OSA. Early detection plays a role in implementing interventions and personalized treatment plans ultimately leading to better patient outcomes and reducing the healthcare burden associated with hypertension linked to OSA.

Our methodology incorporates elements that enhance the strength and effectiveness of the predictive model. By employing timing interval windowing and creating an feature extractor designed specifically for sleep data we successfully captured the intricate temporal relationships present, in polysomnography signals. This method allowed us to extract features that accurately reflect the physiological patterns associated with OSA and hypertension.

Our model combines both clinical data to create a framework for identifying individuals at a higher risk of hypertension within the OSA community.

While our research shows promise there are limitations to be acknowledged. The size of our dataset was relatively small. We faced challenges due to memory constraints during model development and evaluation. To address these issues future studies should focus on validating our model in longitudinal studies with diverse participant groups. Additionally exploring methods for feature extraction and integrating types of data will further improve the accuracy and applicability of our model.

In summary our study highlights the potential of machine learning techniques in enhancing risk prediction and personalized treatment plans for individuals with hypertension related to OSA. By utilizing data from polysomnography tests and applying machine learning approaches our predictive model serves as a resource for healthcare professionals to identify those at risk of hypertension and implement targeted interventions to prevent negative health outcomes. Looking ahead ongoing research efforts and collaboration across disciplines will be crucial in advancing precision medicine strategies for managing hypertension in individuals, with OSA.

## 5 Conclusion

In this research, we investigated methods of extracting features from polysomnography, using the SHHS dataset. Before extracting the features, we prepared the data by removing any distortions and applying a filter to eliminate remaining noise. Additionally, we used timing interval windowing as a method for feature extraction. This allowed us to create a 2D array that made it easier to apply transfer learning techniques. We fine-tuned our model, making it more adaptable to the characteristics of sleep data. Furthermore, we introduced a feature extractor that was specifically designed for this study. This extractor was created with attention to detail to adapt well and accurately represent patterns in polysomnography data. We evaluated our proposed methodology using a ten-fold cross-validation technique to assess the potential generalizability of the model. Our contributions, such as developing feature extraction methods and implementing a tuned adaptive feature extractor, significantly improved the accuracy of predicting incident hypertension in individuals with OSA. When combined with AI and Machine Learning techniques, these new developments in methodology may provide clinicians with tools for early diagnosis and treatment of hypertension in OSA populations. We had a memory limitation due to the high number of features for each patient and the relatively small number of patients in the dataset. Our future research will concentrate on integrating diverse data modalities and exploring innovative feature extraction techniques. Ethical considerations and validation of the novel models in larger longitudinal studies in diverse populations will advance personalized approaches to managing hypertension for individuals with OSA.

## Notes

https://sleepdata.org/datasets/shhs

### Competing Interest Statement

The authors have declared no competing interest.

### Summary of Updates

adds more detailed contextual references to public health efforts, such as clinical applications and impact on healthcare burden, which were not as emphasized in the 2023 version. improvements are discussed in the Word version, indicating refinements made between the two versions.

